# The Body Knows Better: Sensorimotor signals reveal “Suboptimal” inference of the Sense of Agency in the human mind

**DOI:** 10.1101/2024.02.14.579431

**Authors:** Asaf Applebaum, Ophir Netzer, Yonatan Stern, Yair Zvilichovsky, Oz Mashiah, Roy Salomon

## Abstract

Sense of Agency (SoA) is the feeling of control over our actions. SoA has been suggested to arise from both implicit sensorimotor integration as well as higher-level decision processes. SoA is typically measured by collecting participants’ subjective judgments, conflating both implicit and explicit processing. Consequently, the interplay between implicit sensorimotor processing and explicit agency judgments is not well understood. Here, we evaluated in one exploratory and one preregistered experiment (N=60), using a machine learning approach, the relation between a well-known mechanism of implicit sensorimotor adaptation and explicit SoA judgments. Specifically, we examined whether subjective judgments of SoA and sensorimotor conflicts could be inferred from hand kinematics in a sensorimotor task using a virtual hand (VH). In both experiments participants performed a hand movement and viewed a virtual hand making a movement that could either be synchronous with their action or include a parametric temporal delay. After each movement, participants judged whether their actual movement was congruent with the movement they observed. Our results demonstrated that sensorimotor conflicts could be inferred from implicit motor kinematics on a trial by trial basis. Moreover, detection of sensorimotor conflicts from machine learning models of kinematic data provided more accurate classification of sensorimotor congruence than participants’ explicit judgments. These results were replicated in a second, preregistered, experiment. These findings show evidence of diverging implicit and explicit processing for SoA and suggest that the brain holds high-quality information on sensorimotor conflicts that is not fully utilized in the inference of conscious agency.

## Introduction

Volitional, goal-directed actions are one of the most fundamental activities performed by humans. In healthy humans, these movements are accompanied by a constant, fluid, and subtle sense of agency. This Sense of Agency (SoA), is the feeling of control over the body’s motor movements (Gallagher, 2000; Haggard, 2017). SoA is usually unnoticed and considered to have a “thin phenomenology”, thus when unperturbed it resides in the backdrop of our conscious experience rather than being an explicit component of it (Gallagher, 2007; Metzinger, 2004; Tsakiris et al., 2007). According to contemporary theories such as the two-step theory (e.g. Synofzik et al., 2013), SoA comprises two distinct but related phenomena. (1) A pre-reflective *implicit sense of agency* (De Vignemont and Fourneret, 2004; Haggard, 2017; Legrand, 2006), based on integration of low-level sensorimotor cues, and (2) a high-level conceptual *explicit judgments of agency*. Implicit SoA is thought to arise from predictive computations on sensorimotor signals. Motor actions are coupled with an internal efference copy that through internal forward models forms an expectation of the sensory consequences of the action (Von Helmholtz, 1867; Sperry, 1950; von Holst and Mittelstaedt, 1950; Von Holst, 1954; Wolpert et al., 1995; Wolpert and Kawato, 1998). The congruency between the internal prediction and the actual sensory feedback is then computed. If the two are congruent, the feeling of agency over the action is maintained. If the signals are incongruent, the brain will try to compensate for small errors through implicit sensorimotor corrections (Izawa et al., 2008; Izawa and Shadmehr, 2011; Nielsen, 1963; Synofzik et al., 2008a, 2006; Wei and Körding, 2009). However, if the discrepancy is large, the feeling of agency will be disrupted. The two-step theory of agency posits that this implicit process is further modified by higher level expectations, intentions, and beliefs, resulting in an explicit judgment of SoA (Blakemore et al., 2002; Krugwasser et al., 2019; Stern et al., 2020; Harduf et al., 2023; Synofzik et al., 2009; Haggard and Chambon, 2012; Bu-Omer et al., 2021; Le Bars et al., 2020; Wen and Haggard, 2020). The interplay between these implicit sensorimotor processes and explicit metacognitive evaluation of action has been debated (Charles et al., 2020, 2017; Locke et al., 2020; Wen et al., 2023). Most motor actions we perform are implicit and do not require motor awareness (e.g., we don’t have to think about putting one foot in front of the other while casually walking). However, some actions do require conscious performance monitoring that demand cognitive control (e.g., finely chopping vegetables), thus there is also an explicit level in which we can evaluate these actions (Locke et al., 2020). In studies of SoA, the implicit and explicit processes are often difficult to disentangle. Many SoA studies focus mainly on subjective judgments of action control, using explicit agency judgments. These, may likely conflate both low level sensorimotor conflict signals as well as second-level processes, taking into account further higher-level contextual and metacognitive information (Haggard, 2017; Lafleur et al., 2020; Loehr, 2022; Synofzik et al., 2008a). However, recent studies have failed to dissociate the implicit level of sensorimotor control from explicit agency judgments criteria (Chambon et al., 2014; Constant et al., 2022; Wen et al., 2023). For example, Constant et al., found that agency judgments reflect first-order measures of the internal signal, without involving metacognitive computations, suggesting that judgments of agency are simply reports of the implicit level and does not add to the primary sensorimotor information (Constant et al., 2022). It is therefore unclear which information is processed in each level of SoA, and how these two levels interact with one another. It is imperative to understand this relationship in order to truly understand the SoA process.

The dual nature of SoA inference makes its empirical study challenging. A standard method for studying and measuring SoA is to ask participants to make an explicit judgment of their subjective experience of agency or control over an action (Balslev et al., 2007; Chambon et al., 2013; Nahab et al., 2011; Nierula et al., 2021; Sato, 2009; Serino et al., 2022; Stern et al., 2020) or regarding the congruency between their actions and consequences (Farrer et al., 2003; Franck et al., 2001; Krugwasser et al., 2022, 2019; Stern et al., 2021). While this explicit method is simple and direct, it only permits access to the end outcome of the SoA process, possibly conflating the implicit processing of sensorimotor conflicts and higher levels judgments. It has been also pointed out that explicit measures of agency can be subject to response bias, such that, for example, the presence of other agents can alter SoA (Wegner, 2003). Consequently, there have been efforts to develop implicit measures to directly access the implicit component of SoA. The most frequently used implicit measure is the “temporal binding’ approach in which the temporal relation between actions and their consequences is suggested to reflect a feeling of authorship over the action (Haggard et al., 2002; Tsakiris and Haggard, 2003; Moore and Haggard, 2008; Buehner and Humphreys, 2009; Moore and Obhi, 2012; Caspar et al., 2015). One proposal is that temporal binding comes about through an association of forward motor commands with specific, predictable action-effects (Haggard and Clark, 2003), thus, may serve as a good candidate for the implicit SoA component. However, the relationship between temporal binding and SoA has been debated (Dewey and Knoblich, 2014; Saito et al., 2015; Schwarz et al., 2019). For example, Dewey and Knoblich (2014) evaluated explicit agency ratings against temporal binding scores and found no correlation between these measures (Dewey and Knoblich, 2014). In addition, temporal binding is a non-embodied measure. This limits its implications for embodied SoA, which is proposed to have unique characteristics and underlying processing (Stern et al., 2020; Wen et al., 2023). Namely, embodied SoA, focuses on the immediate link between the intention to perform the action and the action itself (e.g., moving your hand), while non-embodied SoA focuses on the link between the action performed by the agent (e.g., moving your finger) and the intended outcome (e.g., pressing a button) (Stern et al., 2020). These results and other similar findings (e.g., Saito et al., 2015; Schwarz et al., 2019) suggest that temporal binding may not be a reliable proxy for SoA. How to investigate the early, implicit stages of SoA processing remains an open question. An appropriate marker for the pre-reflective implicit SoA must tap into the underlying process of it, namely, the detection of sensorimotor conflicts. This may also help to reliably elucidate the relationship between implicit sensorimotor signals and explicit SoA judgments.

A well-studied aspect of implicit sensorimotor control is online adaptation in face of sensorimotor conflicts. For example, in a seminal paper, Nielsen (1963) created a discrepancy between visual and motor-proprioceptive information. When small sensorimotor conflicts were introduced, participants did not report loss of control over their movement, but, unbeknownst to them, made adjustments to their movement to compensate for the discrepancies in sensorimotor cues (Nielsen, 1963). These online sensorimotor corrections were present also when sensorimotor conflicts reached a larger magnitude, but then, participants experienced loss of SoA. Yet, it is unknown how these implicit error signals are incorporated in explicit judgments of SoA.

Prominent theoretical and empirical work have demonstrated that in order to cope with noise in sensorimotor systems the brain uses internal forward models to guide actions (van Beers et al., 2002; Franklin and Wolpert, 2011). Specifically, when executing a motor action, there are often mismatches between the moving limb’s desired and actual position (Franklin and Wolpert, 2011). In order to get from the current estimated state of an action to its desired state, the forward model and sensory feedback are compared and the error between them is calculated (Wolpert et al., 1995; Wolpert and Kawato, 1998). The required motor corrections are then calculated based on the error, resulting in adjustments to the movement online (Wolpert and Ghahramani, 2000; Tseng et al., 2007; Adams et al., 2013). These motor adaptations are typically unnoticed, especially when introducing minor sensorimotor conflicts (Nielsen, 1963; Fourneret and Jeannerod, 1998; Kannape et al., 2010; Salomon et al., 2021; Pereira et al., 2023) which may make them a good candidate for an implicit measurement of the brain’s sensitivity to sensorimotor conflicts and the early stages of SoA processing.

Here, we investigated the relationship between implicit, low-level sensorimotor processing and explicit, higher-level subjective agency judgments. We examined if the sensorimotor conflict information utilized in the implicit level of SoA processing is also used for the explicit level of SoA judgments. To this end, a, data-driven machine learning approach has been applied in this study. We collected two datasets, an exploration dataset (experiment 1) and a replication dataset (experiment 2). For both, we conducted an embodied sensorimotor conflict detection task (Krugwasser et al., 2022, 2019; Stern et al., 2020) in immersive virtual reality, in which participants performed a reaching movement and viewed a virtual hand either performing the same movement or an altered movement. To assess explicit judgments of agency, after each movement participants judged whether their movement was congruent with the movement they observed. Throughout both experiments, participants’ hand kinematics were monitored. In order to evaluate the relationship between implicit and explicit SoA processes at a single trial level, we used two machine learning algorithms to build a classification model for each participant that predicts, based on the participants’ hand kinematics, (1) sensorimotor conflicts and (2) the subjective judgments of agency. We envisioned three possible outcomes (Fig 1C): If implicit kinematics information and explicit judgments of SoA showed similar sensitivity this would suggest that the information from sensorimotor signals is used to inform explicit SoA judgments. If explicit SoA judgments showed higher accuracy than the classification model based on implicit sensorimotor signals, this would suggest that subjective SoA judgments provide additional information enabling superior performance. Finally, as per our preregistered hypothesis, if classification based on implicit sensorimotor information outperformed explicit judgments made by the participants this would suggest that explicit SoA judgments are based on a suboptimal inference of sensorimotor signals.

**Figure 1.**
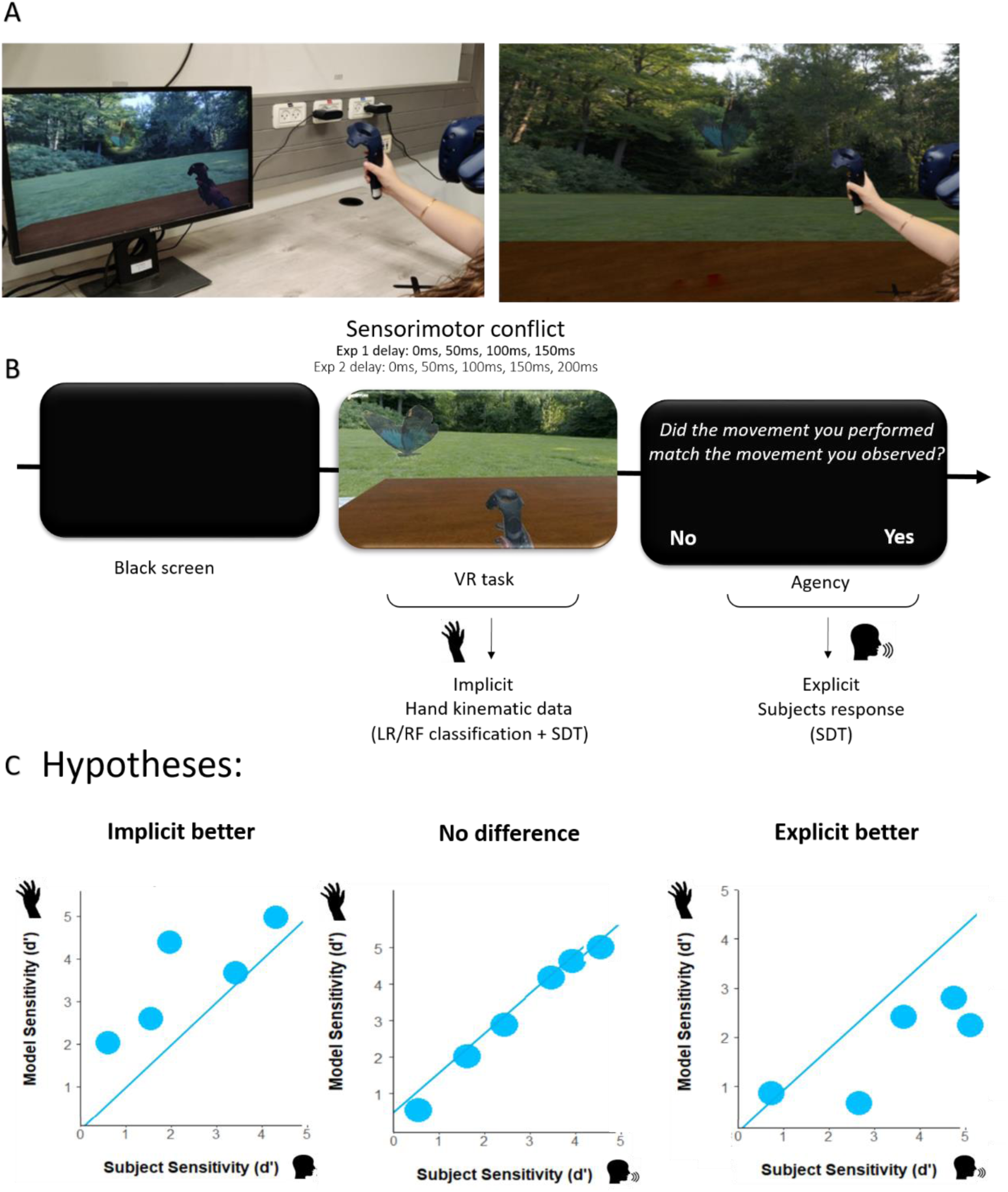
Experimental setup, procedure and hypotheses. (A) Experimental setup. Left: A participant in the experimental setup. The participant is wearing a HMD and using the controller to perform the VH task. The controller is placed on a black ‘x’ sign that marks the start and end locations of the movement in each trial. Right: A conceptual illustration of the participant’s view performing the VH task within the virtual environment. (B) Procedure. Each trial started with a black screen, followed by the presentation of the virtual environment. In the virtual environment, participants controlled a virtual hand (VH). A butterfly appeared in the environment after 200ms. Participants were instructed to “grab” the butterfly. Following this, the participants judged whether the movement they saw was congruent with the movement they made. (C) Hypotheses. Conceptualization of results under each one of the possible hypotheses under our analysis plan. Left: the model based kinematic data will hold more information about the sensorimotor conflict compared with the subjects’ explicit judgments. Middle: both measures will hold the same amount of information. Right: Subjects’ explicit judgments will hold more information about the conflict compared with the kinematic data.

## Methods

Two datasets were collected: an exploration dataset (experiment 1) and a replication dataset (experiment 2). The exploration dataset was utilized to construct analyses pipelines and formulate hypotheses. The selected analysis process and hypotheses were pre-registered before the analysis of the replication dataset (https://aspredicted.org/~lAuCCCl1t6). As explained in the subsequent sections, these two datasets were collected using the same paradigm, with minor modifications.

### Experiment 1

#### Participants

Twenty-five healthy (mean age: 26.5 years, STD: 6.84 years, 14 females) right-handed participants who were unaware of the experiment’s goal participated in the study. None of the participants self-reported a history of mental or neurological impairments, and all had normal or corrected-to-normal vision. All participants gave written informed consent and were compensated (50 NIS per hour or course credits) for their participation. All procedures were performed in compliance with the institutional guidelines of the Gonda Multidisciplinary Brain Research Center and have been approved by the institutional committee (approval number ISU202107003).

Two participants were excluded from all analyses (one for not finishing the experiment and one for failing to have sufficient data, see supplementary methods), resulting in a final dataset of 23 participants.

#### Setup

Participants were seated at a table. They wore the Vive HMD and held the Vive controller in their right hand. They placed the controller on a black ‘x’ sign, a position that marked the beginning and end of each trial’s movement. Participants were instructed to position their left hand underneath the table to preserve the feeling of immersion, as only their right hand’s avatar was displayed (Figure 1A).

#### Virtual Environment

Unity (2018.3.2) was utilized to create a realistic virtual environment. In the virtual setting, participants sat at a table with a black ‘x’ sign marked on it. The virtual table and x sign were positioned in corresponding locations to the table and x sign in the experimental room, to provide matching visual and tactile information. They controlled a virtual hand (VH) that looked and moved similarly to their real hand in space and time. A butterfly was hovering above the table, some distance away from the ‘x’ sign (Figure 1A).

#### Experimental Procedure

Each experiment consisted of a short training session followed by the test trials. Each trial (Figure 1B) started with a black screen (600ms), followed by the presentation of the virtual environment. Following a 200ms interval, the butterfly appeared in one of five predetermined spots (see supplementary methods). Participants were instructed to reach their right hand holding the controller towards the butterfly’s location. During the movement, the butterfly disappeared once the hand moved a certain distance towards it, such that participants could not use it as a feedback cue. In 25% of trials, the VH movement was spatially and temporally congruent with the participant’s hand (congruent condition). A sensorimotor conflict was introduced in the remaining 75% of trials (incongruent condition), either in space (spatial manipulation, angular deviation of the reaching movement to the left) or in time (temporal manipulation, delay between the actual and virtual hand movements). In each manipulation domain, there were four magnitudes of manipulation (0/50/100/150 ms in the temporal domain and 0°/12°/16°/20° in the spatial domain), resulting in different levels of difficulty for the task. Following the reaching movement, to measure the explicit judgments component of SoA, participants responded to a two-alternative forced-choice question (the question was presented in Hebrew): “was the movement I saw congruent with the movement I have made in space and time?” (Franck et al., 2001; Krugwasser et al., 2019; Stern et al., 2020). After the experiment, participants filled out three questionnaires. PQ-B to assess psychotic-like symptoms (Loewy et al., 2005, 2011), IPASE to assess psychotic-like anomalous self-experiences (Cicero et al., 2017), and OCI-R to assess symptoms of obsessive-compulsive disorder (Foa et al., 2002). Participants who did not understand the task or did not complete the experiment were excluded from the analyses.

#### Analysis Plan

This study aimed to evaluate how well explicit SoA judgments and implicit SoA based on sensorimotor conflicts may be inferred from kinematic data and to describe the nature of their relationship. In this study, only temporal sensorimotor conflicts were taken into account. To achieve these objectives, binary classifiers were trained to predict, based on motion kinematics: (1) whether sensorimotor conflict was introduced in the trial and (2) the subjective judgment.

Logistic regression and random forest methods were chosen as classifier algorithms based on their interpretability. Considered a “transparent model”, logistic regression provides a straightforward feature importance analysis. Random forest models are more “opaque”, yet they are still reasonably simple to interpret and have more predictive power (Belle and Papantonis, 2021; Caruana and Niculescu-Mizil, 2006) (Belle and Papantonis, 2021; Caruana and Niculescu-Mizil, 2006).

#### Data Acquisition and Preprocessing

The HTC Vive system tracks (at a frequency of 90Hz) the 3D locations of the participants’ hands throughout the experiment, resulting in three timeseries (location of the X, Y, and Z axes). The first and second derivatives (representing velocity and acceleration, respectively) were computed based on the location timeseries. To correspond with our classification model, the timeseries were transformed into features using ‘tsfresh’, a python package specialized in extracting features from timeseries (Christ et al., 2018). Features with zero variance across all participants were eliminated, resulting in 98 features per trial, which construct the full feature representation (see supplementary for more details). An algorithm based on the Variation Inflation Factor (VIF) was used to eliminate multi-collinearity (Yoo et al., 2014), resulting in a subset of 17 relatively independent features which construct the clean feature representation. (See pseudo-code and more details on the features in the supplementary). Trials in which participants gave no response or made invalid movements were removed from the analysis. Participants that did not have enough trials in each condition were excluded from the analysis (see Supplementary methods for exclusion criteria).

#### Model Evaluation

There were two categories of classification: predicting the trial’s condition (whether sensorimotor conflict was introduced), or an objective classification, and predicting the subjective judgment, or a subjective classification. Each category was divided into a few sub-classifications (see supplementary methods). Classifiers were trained, using the full feature representation, for each classification category in each participant separately. A 10-fold cross-validation method was used for the classification category to evaluate classifier performance. Since the datasets for most classifications were imbalanced, AUC was employed as an evaluation metric.

To determine the degree to which participants’ kinematic signatures are similar, the similarity between each participant’s logistic regression models was calculated using the Pearson correlation between the coefficients of these models. Similarly, we examined the similarity of participants’ behavior by cross classification. The cross-subject model technique included training classifiers iteratively on the data of a single participant, evaluating it on the data of the remaining participants.

To understand the effect of each feature on the classification models, both logistic regression and random forest feature importance analyses were extracted. In logistic regression models, the regression coefficients represent the weighted contribution of each feature to the model, thus they determine the importance of the features. The importance of the features in the random forest models was determined by calculating the mean decrease in impurity for each feature (Molnar, 2022).

Participants’ performance was measured by quantifying the sensitivity and bias (Swets, 1964; Stanislaw and Todorov, 1999; Stern et al., 2020; Krugwasser et al., 2022; Macmillan, 2002; Green and Swets, 1966) of signal detection theory (SDT). Classifiers’ sensitivity and bias were also calculated to compare the performance of the classifiers to that of the participants.

Pearson correlations were used to assess the relationship between model performance, participants’ SDT performance and questionnaire ratings. Multiple comparisons were accounted for using the Bonferroni method.

#### Hardware

The experiment was conducted on a laptop (CPU: Intel Core i9 processor, RAM: 32 GB, GPU: NVIDIA GeForce RTX 2070) running in-house software (developed using Unity 2018.3.2). An HTC Vive HMD, a Vive controller, and two base stations were used to track participants’ hand motions and generate the virtual environment projected through the HMD (refresh rate 90HZ). The trackpad of the Vive controller was employed as an input device to record participants’ answers (Figure 1A).

#### Hypotheses

Based on our exploratory dataset, we pre-registered our hypotheses and analyses pipeline (https://aspredicted.org/~lAuCCCl1t6). We hypothesized that:

H_1_: Our models would be able to predict sensorimotor conflicts from the hand motor kinematics.

H_2_: Kinematic-based models of different participants would be similar, meaning that the relationship between hand motor kinematics and sensorimotor conflicts would also be similar across participants.

H_3_: Models’ sensitivity to these conflicts would be greater than the participants’ sensitivity.

H_4_: Our model would also predict subjective judgment, however, at a lower accuracy than sensorimotor conflicts.

By comparing subjects’ sensitivity to the sensitivity of their models that were based on kinematic data, we tried to understand which information is used in each step of the SoA process (explicit vs. implicit). Figure 1C presents the three possible outcomes of this comparison. That is, if models’ sensitivity to sensorimotor conflicts is greater than subjects’ sensitivity, then it can be interpreted as that some information in the implicit level of SoA is disregarded when moving to the higher, explicit level of SoA (Figure 1C, left panel). If subjects’ sensitivity to the task is greater than their model’s sensitivity, it means that additional information is integrated in the explicit level of SoA processing (Figure 1C, right panel). The third alternative is that both steps hold the same information (Figure 1C, middle panel).

### Experiment 2

As stated above, experiment 2 was a pre-registered replication study of experiment 1. Unless otherwise specified, Experiment 2 was identical to Experiment 1.

#### Participants

Thirty-five healthy (mean age: 24.8 years, STD: 4.46 years, 28 females), which met the same criteria as experiment 1, participated in the study. Three participants were excluded for failing to satisfy the trials threshold after filtering, resulting in 32 participants.

#### Experimental Procedure

The experimental procedure was identical to that of experiment 1 except for the following modifications: This experiment included only manipulations in the temporal domain. The magnitudes of the temporal manipulations were 0, 50, 100, 150, and 200 milliseconds. This experiment’s ‘test’ phase consisted of 256 trials, 64 congruent trials, and 48 incongruent trials for each magnitude. These trials were separated into four blocks of 40 and two blocks of 48. There were six predetermined locations in which the butterfly could appear (see supplementary).

## Results

### Experiment 1

As expected and in line with previous research (e.g., Krugwasser et al., 2019; Stern et al., 2020), participants showed a gradual and significant decline in SoA as the degree of delay increased (𝐹(3,66) = 110.4, 𝑝 < 0.00001, 𝜂_p_^2^ = 0.834) (Figure 2A).

**Figure 2.**
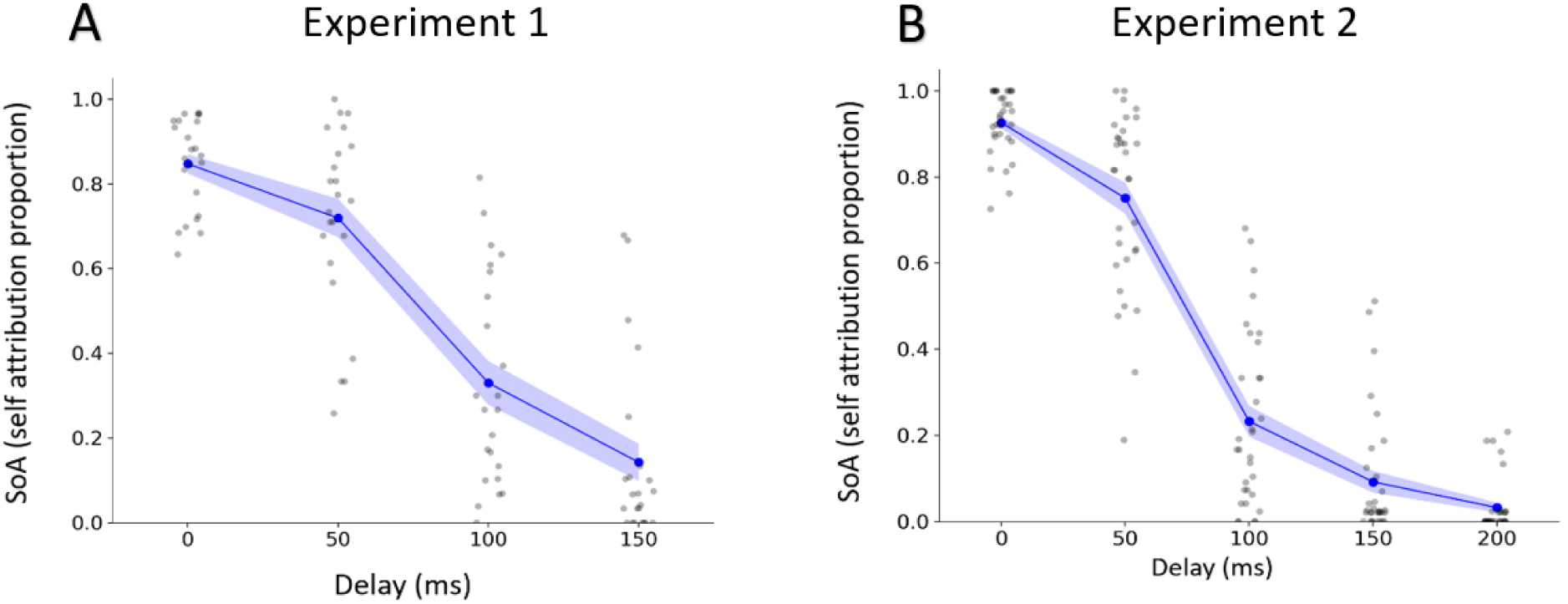
Participants’ self-attribution proportion with respect to manipulation magnitude. Blue dot represents the group mean for each manipulation magnitude, blue ribbon represents the SE. Grey dots represent individual participants. (A) Participants’ performance in experiment 1 (SoA_0_=0.85, SoA_50_=0.72, SoA_100_=0.33, SoA_150_=0.14). (B) Participants’ performance in Experiment 2 (SoA_0_=0.92, SoA_50_=0.75, SoA_100_=0.23, SoA_150_=0.09, SoA_200_=0.03).

Classifiers trained on kinematic data (AUC_LR_ = 0.96±0.04, AUC_RF_ = 0.93±0.06) showed significantly high levels of differentiation between congruent and incongruent trials (𝑡_𝐿𝑅_(22) = 47.51, 𝑡_𝑅𝐹_(22) = 34.54, 𝑝 < 0.00001 for both). The logistic regression model outperformed the random forest (𝑡(22) = 5.22, 𝑝 < 0.0001). Both classifiers had significant performance differences surviving Bonferroni correction between the 50ms and the 100ms conditions (𝑡_𝐿𝑅_(22) = 4.76, 𝑡_𝑅𝐹_(22) = 6.52, 𝑝 < 0.001 for both), but not between the 100ms and the 150ms conditions, as AUC levels in these two highest levels of alterations were both extremely high and above 0.95 (Figure 3A&B). To directly compare participants’ performance based on explicit SoA judgments and the classification models’ performance based on kinematic data, we calculated SDT measurements. The sensitivity of both classifier models (𝑑^′^ = 2.82 ± 0.83, 𝑑^′^ = 2.37 ± 0.71) was determined and compared to participants’ sensitivity (𝑑^′^ = 1.13 ± 0.54). Both classifier models outperformed participants (𝑡_𝐿𝑅_(22) = 8.97, 𝑡_𝑅𝐹_(22) = 7.34, 𝑝 < 0.00001 for both) (Figure 3C&D). To further compare explicit and implicit SoA, we evaluated models’ performance in classifying participants’ explicit SoA judgments based on kinematic data. These models (AUC_LR_ = 0.76±0.09, AUC_RF_ = 0.78±0.08) showed a significant ability to differentiate between participants’ explicit SoA judgments (me vs. not me) trials (𝑡_𝐿𝑅_(22) = 14.02, 𝑡_𝑅𝐹_(22) = 15.62, 𝑝 < 0.00001 for both). However, both classifiers performed better on the objective than the subjective classification categories (𝑡_𝐿𝑅_(22) = 11.13, 𝑡_𝑅𝐹_(22) = 9.12, 𝑝 < 0.00001 for both). Since the correlation of classifiers’ performance between the LR and the RF models was high, positive, and statistically significant (𝑟(21) = 0.96, 𝑝 < 0.00001), we further focused on the LR model due to its high interpretability. To evaluate the contribution of each feature to the classification models, the feature importance analysis was calculated and revealed similarities between the two classifier methods. The three most important features related to velocity and acceleration of the hand movement (mean vertical velocity, total acceleration in the vertical axis and mean acceleration in the depth axis) and were the same in both classifiers (Figure 3E&F). Examining the similarity of the logistic regression model between participants revealed a high degree of similarity (0.72±0.11) between participants’, suggesting that participants implicit motor behaviors were similar (Figure 3G). To assess the relationship between implicit SoA processes and tendencies for abnormal perceptual and bodily experiences, Pearson correlations were calculated between model performance and participants’ questionnaire ratings. As expected, positive and significant correlations were found between all questionnaire scores. No significant correlations were found between questionnaire scores and the remaining variables (Figure 3H).

**Figure 3.**
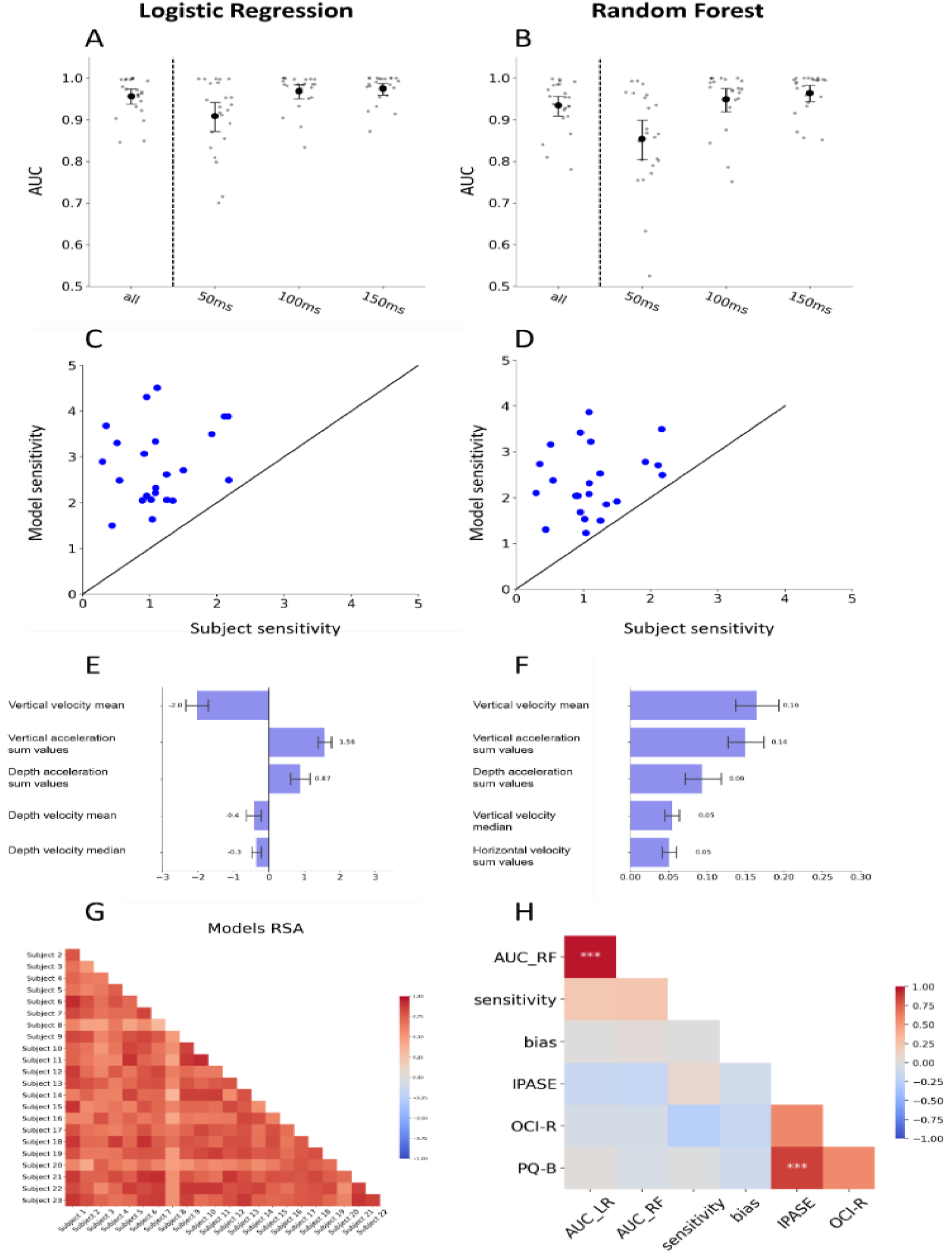
Classifier results of experiment 1: objective classification. A,B average AUC across participants for each objective classification separately. The large black dot represents the group mean, the error bars represent the bootstrapped 95% confidence interval, and the small grey dots represent individual participants. (A) Logistic regression classifier (mean_all_=0.96, mean_50_=0.91, mean_100_=0.97, mean_150_=0.975). (B) Random forest classifier (mean_all_=0.93, mean_50_=0.85, mean_100_=0.95, mean_150_=0.96). C, D comparison between models’ and participants’ sensitivity (d’). (C) Logistic regression. (D) Random forest. E,F results of feature importance analysis in both models. (E) Feature importance in the logistic regression as determined by the average beta score of each feature, and (F) feature importance in the random forest as determined by the mean decrease in impurity approach. (G) Correlation between betas in different participants’ logistic regression models (mean_r_=0.72, STD=0.12). (H) Pearson correlation between the performance of logistic regression, random forest, participants d’ and bias, and questionnaire scores. *p < 0.05 **p < 0.01 ***p < 0.001.

### Experiment 2

In line with the results of experiment 1 and the pre-registration, participants’ responses to the agency question show a gradual and significant decline in SoA as the degree of delay increased (𝐹(4,124) = 368.484, 𝑝 < 0.00001, 𝜂_p_^2^ = 0.922) (Figure 2B). In addition, both classifiers (AUC_LR_ = 0.97±0.04, AUC_RF_ = 0.96±0.05) were able to differentiate significantly above chance between congruent and incongruent trials (𝑡_𝐿𝑅_(31) = 72.27, 𝑡_𝑅𝐹_(31) = 55.41, 𝑝 < 0.00001 for both). The logistic regression model outperformed the random forest (𝑡(31) = 5.47, 𝑝 < 0.0001). Both classifiers significantly differed in performance between the 50ms and the 100ms classifications (𝑡_𝐿𝑅_(31) = 4.04, 𝑡_𝑅𝐹_(31) = 6.58, 𝑝 < 0.001 for both) surviving Bonferroni correction. Random forest classifiers significantly differed in performance between the 100ms and the 150ms classifications (𝑡_𝑅𝐹_(31) = 3.91, 𝑝 < 0.001) and between the 150ms and the 200ms classifications (𝑡_𝑅𝐹_(31) = 3.33, 𝑝 < 0.01). However, the logistic regression classifier did not differ significantly in performance between these two pairs (𝑡_𝐿𝑅_(31) = 2.64, 𝑝 = 0.013; 𝑡_𝐿𝑅_(31) = 1.25, 𝑝 = 0.22 corerspodently) (Figure 4A&B). In contrast to the pre-registration and experiment 1 results, some participants demonstrated greater sensitivity than the classifiers. Nonetheless, both classifiers (𝑑^′^𝐿𝑅 = 3.28 ± 0.85, 𝑑^′^𝑅𝐹 = 2.77 ± 0.87) outperformed participants (𝑑^′^*_participants_* = 2.27 ± 0.61) at the group level (𝑡_𝐿𝑅_(31) = 5.48, 𝑡_𝑅𝐹_(31) = 2.61, 𝑝 < 0.02 for both) (Figure 4C&D). Additionally, models trained on kinematic data (AUC_LR_ = 0.88±0.05, AUC_RF_ = 0.89±0.05) showed a significant ability to differentiate between participants’ explicit SoA judgments (me vs. not me) trials (𝑡_𝐿𝑅_(31) = 38.55, 𝑡_𝑅𝐹_(31) = 39.07, 𝑝 < 0.00001 for both). However, and in line with the results of experiment 1, both classifiers performed better on the objective than the subjective classification categories (𝑡_𝐿𝑅_(31) = 9.47, 𝑡_𝑅𝐹_(31) = 7.93, 𝑝 < 0.00001 for both). The correlation of classifiers’ performance between the LR and the RF models was again high, positive, and statistically significant (𝑟(30) = 94, 𝑝 < 0.00001) and the feature importance analysis revealed similarities between the two classifiers. As in experiment 1, the top two features related to velocity and acceleration of the hand movement (total acceleration in the vertical axis and total acceleration in the horizontal axis) and were the same in both classifiers (Figure 4E&F). In line with experiment 1, logistic regression coefficients were similar among participants (0.76 ± 0.11) (Figure 4G). As expected, positive and significant correlations were found between all questionnaire scores (Figure 4H). However, no significant correlations were found between questionnaires scores and the remaining variables (Figure 4H).

**Figure 4.**
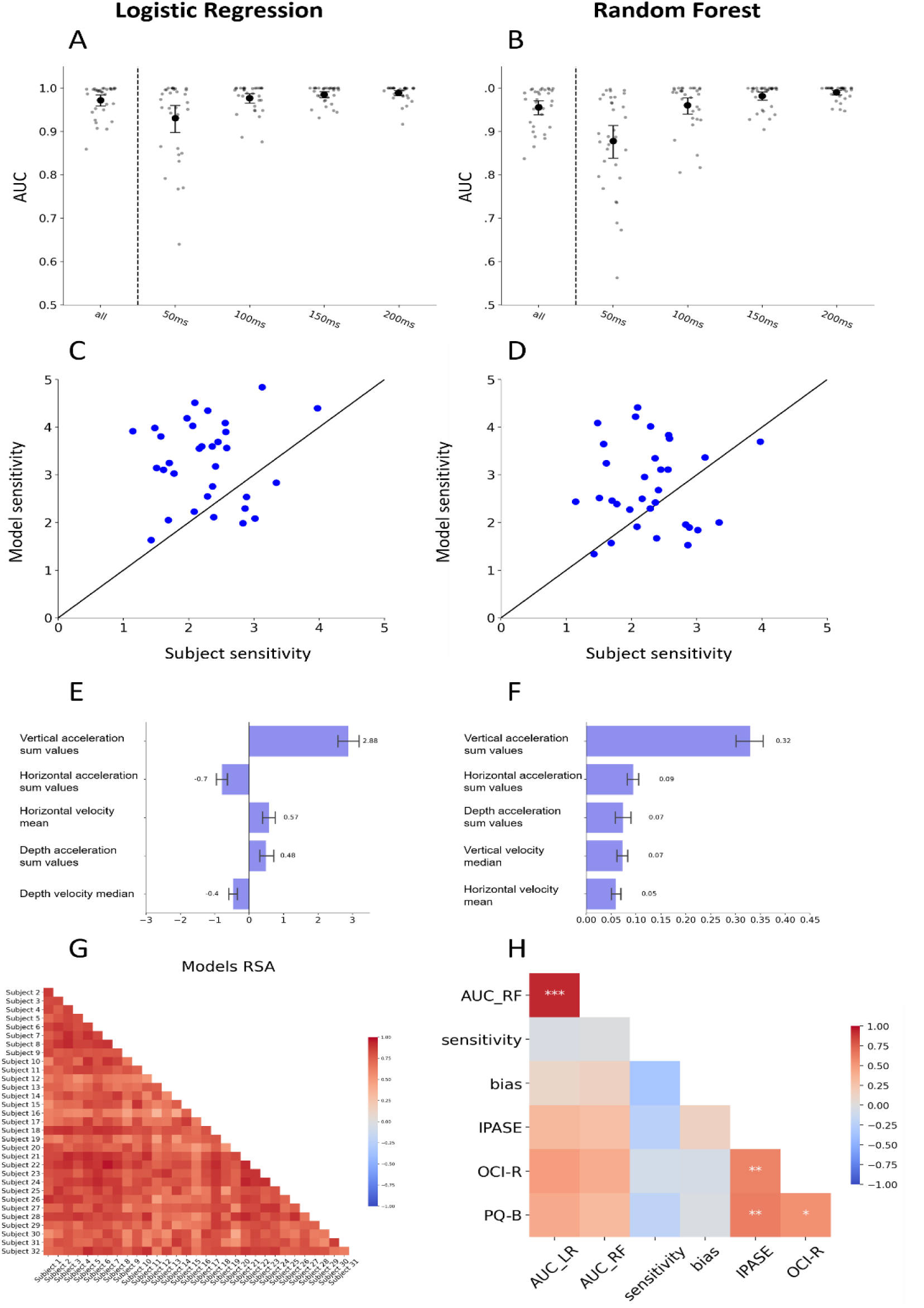
Classifier results of experiment 2: objective classification. A,B average AUC across participants for each objective classification separately. The large black dot represents the group mean, the error bars represent the bootstrapped 95% confidence interval, and the small grey dots represent the individual participants. (A) Logistic regression classifier (mean_all_=0.97, mean_50_=0.93, mean_100_=0.98, mean_150_=0.985, mean_200_=0.99). (B) Random forest classifier (mean_all_=0.95, mean_50_=0.87, mean_100_=0.96, mean_150_=0.98, mean_200_=0.99). C, D comparison between models’ and participants’ sensitivity (d’). (C) Logistic regression. (D) Random forest. E,F results of feature importance analysis in both models (E) Feature importance in the logistic regression as determined by the average beta score of each feature, and (F) feature importance in the random forest as determined by the mean decrease in impurity approach. (G) Correlation between betas in different participants’ logistic regression models (mean_r_=0.76, STD=0.11). (H) Pearson correlation between the performance of logistic regression, random forest, participants d’ and bias, and questionnaire scores. *p < 0.05 **p < 0.01 ***p < 0.001

## Discussion

In this pre-registered study, we investigated the relationship between sensorimotor conflicts, judgments of agency, and hand kinematics using data-driven machine-learning techniques with a novel VR paradigm. Our results, demonstrated in experiment 1 and replicated in experiment 2, reveal several important findings. First, our data-driven models could predict with high accuracy objective sensorimotor conflicts from hand kinematics (H_1_), thereby providing a novel method to implicitly measure the processing of sensorimotor conflicts on a trial by trial basis. In addition, the models were similar across participants (H_2_), indicating that the motor signature of these sensorimotor conflicts is robust and can be generalized to other individuals. Second, hand motor kinematics-based models demonstrated greater sensitivity to sensorimotor conflicts than the participants themselves (H_3_), suggesting that the processes underlying explicit agency judgments are suboptimal in comparison to the information held by the brain regarding sensorimotor conflicts. Finally, our models could predict subjective agency judgments from hand kinematics, but to a lesser extent than sensorimotor conflicts (H_4_).

We found that sensorimotor conflicts could be detected with near-perfect accuracy from hand motor kinematics. This finding is consistent with a large body of evidence from multiple studies demonstrating that sensorimotor conflicts alter motor kinematics to minimize prediction error, frequently without awareness (Nielsen, 1963; Fourneret and Jeannerod, 1998; Fourneret et al., 2001; Knoblich and Kircher, 2004; Kannape et al., 2010; Izawa and Shadmehr, 2011; Oh and Schweighofer, 2019; Marko et al., 2012; Wei and Körding, 2009). These studies showed that when participants experience a spatial discrepancy between their actual and perceived movement, they automatically and, for the most part, unknowingly compensate for that discrepancy (Nielsen, 1963). Here, we extend these findings by revealing that sensorimotor conflicts affect motor kinematics at the single trial level and are similar across participants as indicated by cross-participants decoding. Moreover, to the best of our knowledge, this is the first study to demonstrate the relationship between online motor kinematics and sensorimotor conflicts in the temporal domain.

The novel approach employed in this study enabled us to shed new light on implicit agency processing phases. It is common practice to study agency by measuring subjective sensitivity to sensorimotor conflicts (Franck et al., 2001; Leube et al., 2003; Farrer et al., 2008; Limanowski et al., 2017; Nahab et al., 2011; Krugwasser et al., 2019; Stern et al., 2020; Krugwasser et al., 2021). By predicting sensorimotor conflicts from hand kinematics, we can implicitly measure the brain’s sensitivity to these conflicts without biases of other cognitive processes (e.g., expectations).

The classification models based on motor kinematics demonstrated higher sensitivity in detecting sensorimotor conflicts than the participants themselves. This finding may provide support for the two-step model of agency (Synofzik et al., 2008a, 2013) as it demonstrates that conflict detection is distinct from conscious, explicit judgment processes. Furthermore, it indicates that motor kinematics contain better information about sensorimotor conflict than what is used to form participants’ conscious awareness of these conflicts. This finding suggests that although the brain’s motor system contains high-fidelity information on sensorimotor conflicts, it is not fully utilized in the inference of explicit agency. Theoretically, it appears that the brain processes agency in a suboptimal manner given the information it possesses.

One possible explanation for this phenomenon is that the brain deliberately ignores certain information to generate a unified experience of the self. The continuity of self-awareness and agency is self-evident; healthy humans do not experience frequent disturbances in their sense of agency. Rather, it is characterized by subtlety and pre-reflection, which are components of its “thin phenomenology” (Metzinger, 2004; Tsakiris et al., 2007; Gallagher, 2007). It is also well-established that sensorimotor integration processes modulate sense of agency (Daprati et al., 1997; Frith et al., 2000; Franck et al., 2001; Blakemore et al., 2002; Synofzik et al., 2008a, 2008b, 2013; Krugwasser et al., 2019; Stern et al., 2020). These two properties of sense of agency are challenging to reconcile since both the motor and the sensory systems are prone to neuronal and environmental noise (van Beers et al., 2002; Franklin and Wolpert, 2011). If the brain had utilized all of the information in these systems, the outcome may have been a noisy and unstable experience of agency, resulting in a noisy and unstable self-experience. In order to manage this noise and maintain a consistent sense of agency, the brain may employ a seemingly “suboptimal” inference strategy, not utilizing kinematic error information fully for explicit judgments of SoA. Indeed, participants show wide “tuning curves” for the sense of agency accepting actions with delays of up to 120 ms as their own (Krugwasser et al., 2022, 2019; Stern et al., 2021, 2020). Thus, while task performance for sensorimotor conflicts is suboptimal, SoA and the bodily self are preserved in the face of noisy environments (Zaidel and Salomon, 2023).

Motor kinematics-based models were similar across participants, suggesting that despite motor variability (Latash et al., 2002; Todorov and Jordan, 2002; Dhawale et al., 2017), the motor signature of temporal sensorimotor conflicts is robust and generalizable. Specifically, vertical velocity and acceleration were two of the most predictive features across models. When a sensorimotor conflict was introduced, the mean velocity tended to be lower, but the total acceleration tended to be higher suggesting a form of motor correction (Wolpert and Ghahramani, 2000; Tseng et al., 2007; Adams et al., 2013). When the perceived virtual hand lagged behind the actual hand, the motor system tried to reduce the difference by slowing down the hand’s velocity while keeping it moving toward the target. This process would reduce velocity but increase the number of velocity changes, resulting in a higher total acceleration. This finding is consistent with previous studies indicating that when spatial manipulation is introduced, participants’ movement tends to minimize the difference between the expected and actual position (Nielsen, 1963; Fourneret and Jeannerod, 1998; Kannape et al., 2010; Salomon et al., 2021). This finding suggests, as expected, that the kinematic signature of sensorimotor conflicts stems from online motor adaptations.

This finding may also serve as a basis for a simple and inexpensive diagnostic tool for schizophrenia. Patients with schizophrenia have impaired sensitivity to sensorimotor conflicts (Franck et al., 2001; Krugwasser et al., 2021; Salomon et al., 2021). Future research should evaluate if motor kinematics of schizophrenia patients are also affected by reduced sensitivity, possibly creating a distinctive kinematic signature.

This study has several limitations. First, our models were based on a relatively simple logistic regression method and includes basic features in our models (e.g., mean velocity). These might not be the best for fully describing the explicit and implicit processes of SoA. Moreover, our classification models were based on kinematic data generated from a single effector (right hand) performing a specific trained-on movement. Future studies should attempt to replicate this effect using kinematics of different effectors and more complex movements. Importantly, further research is required to understand if these findings extend to other kinematic systems, such as the oculomotor system.

## Conclusions

The current study investigated the relationship between implicit kinematic processes and explicit judgments of agency. We discovered, using a machine learning approach, that motor adaptations hold information about the early, implicit stages of agency in the form of sensorimotor conflict detection processes. Furthermore, we found that kinematic motor adaptations are more sensitive to sensorimotor conflicts than subjects’ agency judgments. These results demonstrate that the brain retains valuable sensorimotor conflict information for online motor adaptations, that isn’t fully harnessed for the explicit agency inference process. Thus, we have demonstrated that the early and implicit stages of sense of agency can be measured directly from hand kinematics, and that information regarding minor incongruences in the movement is filtered out in the explicit level of agency, perhaps to maintain a smooth and continuous experience of agency and of the self.

## Author contribution

**Asaf Applebaum**: Conceptualization, Investigation, Methodology, Formal analysis, Visualization, Writing – original draft. **Ophir Netzer**: Conceptualization, Investigation, Methodology, Data curation, Writing – review & editing. **Yonatan Stern**: Conceptualization, Investigation, Methodology, Writing – review & editing. **Yair Zvilichovsky**: Software. **Oz Mashiah**: Data curation. **Roy Salomon**: Conceptualization, Methodology, Supervision, Validation, Writing review & editing, Funding acquisition.

## Acknowledgements

This study was supported by the Israeli Science Foundation grant (#1169/17) and a European Union grant (ERC, UNREAL, 949010) to R.S.

## Supplementary

### Methods

#### Setup and virtual environment

Participants were seated at a table. They wore the Vive HMD and held the Vive controller in their right hand. They placed the controller on a black ‘x’ sign that marked the beginning and end of each trial’s movement. The ‘x’ sign was outlined using textured duct tape so that participants could use their touch to find it while wearing the VIVE HMD. Participants were instructed to position their left hand underneath the table to preserve the feeling of immersion, as only their right hand’s avatar was displayed.

In the virtual setting, participants sat at a table with a black ‘x’ sign marked on it. The virtual table and x sign were positioned in corresponding locations to the table and x sign in the experimental room, to provide matching visual and tactile information. They controlled a virtual hand (VH) that looked and moved similarly to their real hand in space and time. A butterfly was hovering above the table, some distance away from the ‘x’ sign (Figure 1A).

#### Experimental Procedure

Each experiment consisted of a short training session followed by the test trials. The purpose of the training was to introduce participants to virtual environment and the task and allow the experimenter to verify that they understood the task correctly. Only test trials were used for data collection. Each trial started with a black screen (600ms), followed by the presentation of the virtual environment. Following a 200ms interval, the butterfly appeared in one of five predetermined spots (see supplementary methods). The participants were instructed to reach their right hand towards the butterfly’s location, after which they were to return to the starting position (marked by an “X” in both real and virtual environments) to end the trial. During the movement, the butterfly would disappear once the hand moved 25 cm from the starting position, such that the participants could not utilize it as a feedback cue. In 25% of trials, the VH movement was spatially and temporally congruent with the participant’s hand (congruent condition). A sensorimotor conflict was introduced in the remaining 75% of trials (incongruent condition), either in space (spatial manipulation, angular deviation of the reaching movement to the left) or in time (temporal manipulation, delay between the actual and virtual hand movements). In each manipulation domain, there were four magnitudes of manipulation (0/50/100/150 ms in the temporal domain and 0°/12°/16°/20° in the spatial domain), resulting in different levels of difficulty for the task. Following the reaching movement, to measure the explicit judgments component of SoA, participants responded to a two-alternative forced-choice question (the question was presented in Hebrew): “was the movement I saw congruent with the movement I have made in space and time?” (Franck et al., 2001; Krugwasser et al., 2019; Stern et al., 2020). Participants were instructed to respond using the Vive controller. The experiment consisted of 240 trials, 30 from each domain’s magnitudes. These trials were divided into six blocks of 40 trials each, with self-paced breaks between them. Within each block, the distribution of trials was identical, but the order of the trials was randomized. After the experiment, participants filled out three questionnaires. PQ-B to assess psychotic-like symptoms (Loewy et al., 2005, 2011), IPASE to assess psychotic-like anomalous self-experiences (Cicero et al., 2017), and OCI-R to assess symptoms of obsessive-compulsive disorder (Foa et al., 2002). Participants who did not understand the task or did not complete the experiment were excluded from the analyses.

#### Butterfly Locations

In the two experiments, butterfly locations were arranged differently. The difference from experiment 1 was due to an additional task performed in conjunction with the second experiment, that requires a change in the butterfly locations.

**Figure S1.**
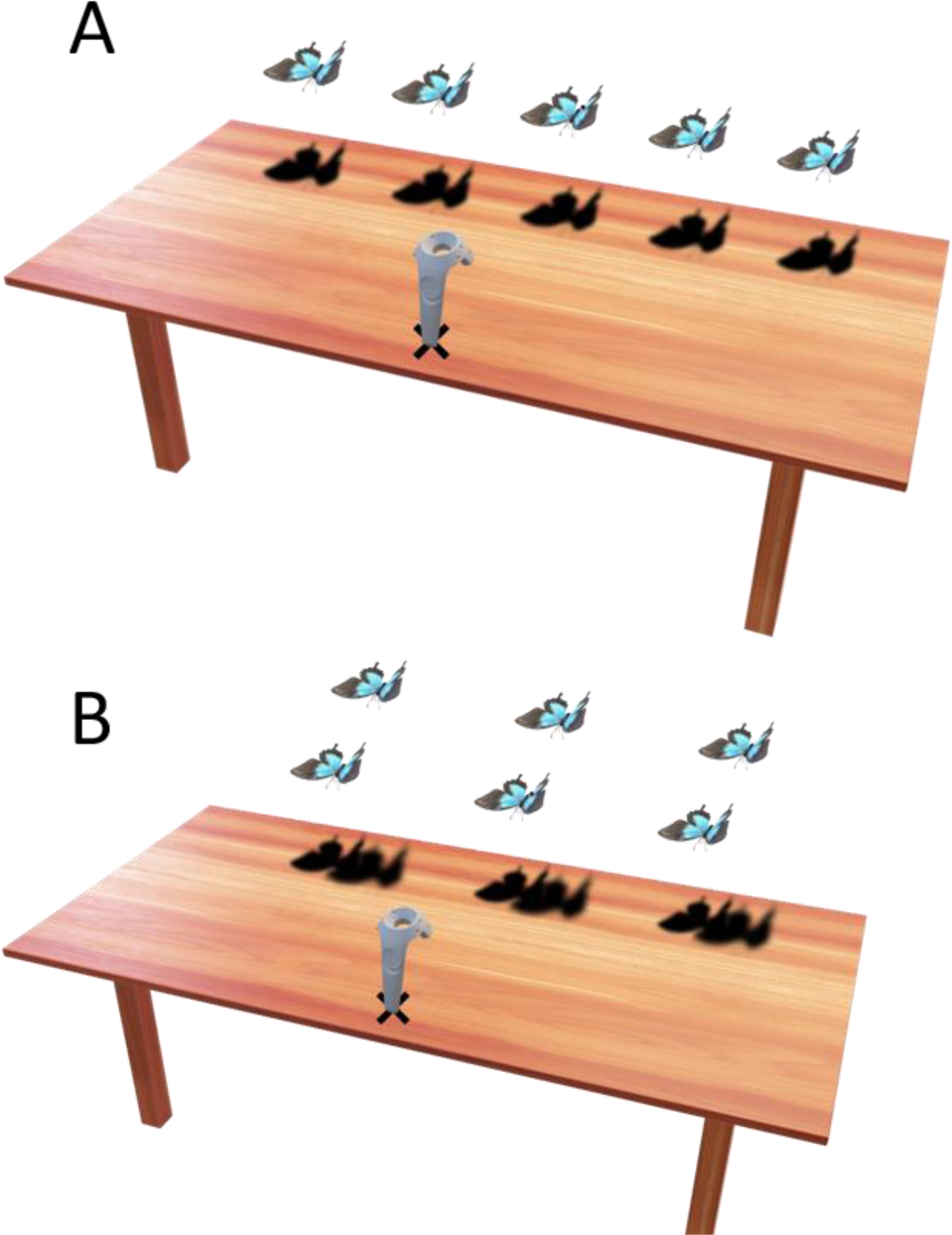
Butterfly positions. (A) Experiment 1 (B) Experiment 2

**Table S1.**
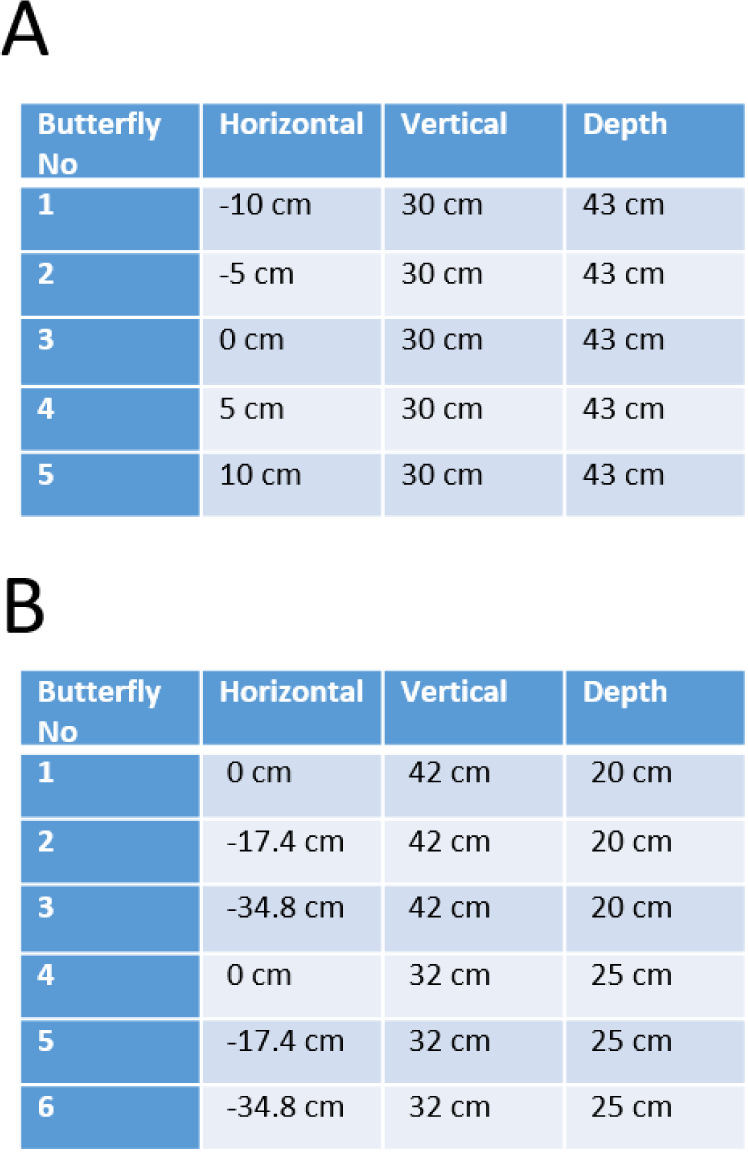
Butterfly positions. All locations are relative to the hand starting point, i.e. the black ‘x’ (A) Experiment 1 (B) Experiment 2

#### Training

The training consisted of 20 trials in which the participants were introduced to all possible levels of alterations / conditions and were given feedback from the experimenter.

#### Data Acquisition and Preprocessing

The HTC system tracks (at a frequency of 90Hz) the 3D locations of the participants’ hands throughout the experiment, resulting in three timeseries (location of the X, Y, and Z axes). The intervals between samples were distributed unevenly. To accurately compare trials and determine velocity and acceleration, each timeseries was interpolated (cubic interpolation) and sampled every 11ms.

##### Filter Trials

Some trials were deleted due to technical difficulties or poor participant performance as part of the data preprocessing. If one of the following conditions was met, the trial was deleted:

- The participant did not answer the question at the end of the trial.
- The duration of the trial was less than 600ms.
- The total depth axis movement was too long (more than 65cm).
- The reaching movement in the trial was too short (less than 10cm from the beginning point to the most distant point).
- The participant remained at the butterfly location for longer than one second.

In experiment 1 Participants who did not have at least 20 trials in each condition were excluded from the analysis following trial elimination. In the remaining participants, 97% of trials were preserved. In experiment 2 the same filters were applied, but in this case, participants who did not have at least 24 trials in each condition were excluded from the analysis following trial elimination. In the remaining participants, 93% of trials were preserved.

##### Velocity and Acceleration Calculation

The first and second derivatives (representing velocity and acceleration, respectively) were computed based on the location timeseries. In addition, total velocity and acceleration timeseries (Euclidean norm of the unidimensional timeseries) were computed, resulting in 11 timeseries for each trial.

##### Feature Representation

The models used in this study require data in feature representation rather than timeseries representation. Therefore, the 11 timeseries were transformed into features using ‘tsfresh’, a python package specialized in extracting features from timeseries (Christ et al., 2018). The following features were calculated from each timeseries: length, maximum value, minimum value, median value, mean value, root mean square, STD, sum of values, and variance, resulting in 99 features per trial.

Features with zero variance across all participants were eliminated, resulting in 98 features per trial, which construct the full feature representation.

##### Multicollinearity Elimination

An algorithm based on the Variation Inflation Factor (VIF) was used to eliminate multi-collinearity (Yoo et al., 2014). In experiment 1, features with high VIF scores were dropped from the dataset resulting in a subset of 17 relatively independent features which construct the clean feature representation. In experiment, 2 the multi-collinearity elimination algorithm has created a subset of 15 relatively independent features which constructed the clean feature representation (see pseudo-code and more details on the features below).

#### Features

The minimum of the total acceleration was discarded in both experiments 1 and 2 due to zero variance. The features that which construct the clean feature representation are detailed in the table below:

**Table S2.** Clean feature representation. A subset of relatively independent features is selected through the VIF-based feature elimination algorithm.

| Feature number | Experiment 1 | Experiment 2 |
| --- | --- | --- |
| 1 | Horizontal location minimum | Horizontal location minimum |
| 2 | Horizontal location variance | Depth location maximum |
| 3 | Depth location sum of values | Horizontal velocity mean |
| 4 | Horizontal velocity median | Horizontal velocity median |
| 5 | Horizontal velocity sum of values | Vertical velocity median |
| 6 | Vertical velocity mean | Depth velocity median |
| 7 | Vertical velocity median | Depth velocity sum of values |
| 8 | Depth velocity median | Horizontal acceleration median |
| 9 | Depth velocity sum of values | Horizontal acceleration sum of values |
| 10 | Horizontal acceleration median | Vertical acceleration median |
| 11 | Horizontal acceleration sum of values | Vertical acceleration minimum |
| 12 | Vertical acceleration median | Vertical acceleration sum of values |
| 13 | Vertical acceleration minimum | Depth acceleration median |
| 14 | Vertical acceleration sum of values | Depth acceleration sum of values |
| 15 | Depth acceleration median | Total velocity minimum |
| 16 | Depth acceleration sum of values |  |
| 17 | Total velocity minimum |  |

#### VIF Algorithm Pseudocode

In each iteration, the VIF score of each feature across participants was calculated, resulting in a vector of VIF scores per feature in the initial stage. In the subsequent phase, only the minimum VIF score for each feature was considered. Then, the feature with the highest minimum VIF score was eliminated. This procedure ended when the highest minimal VIF score fell below five.

**Figure S2.**
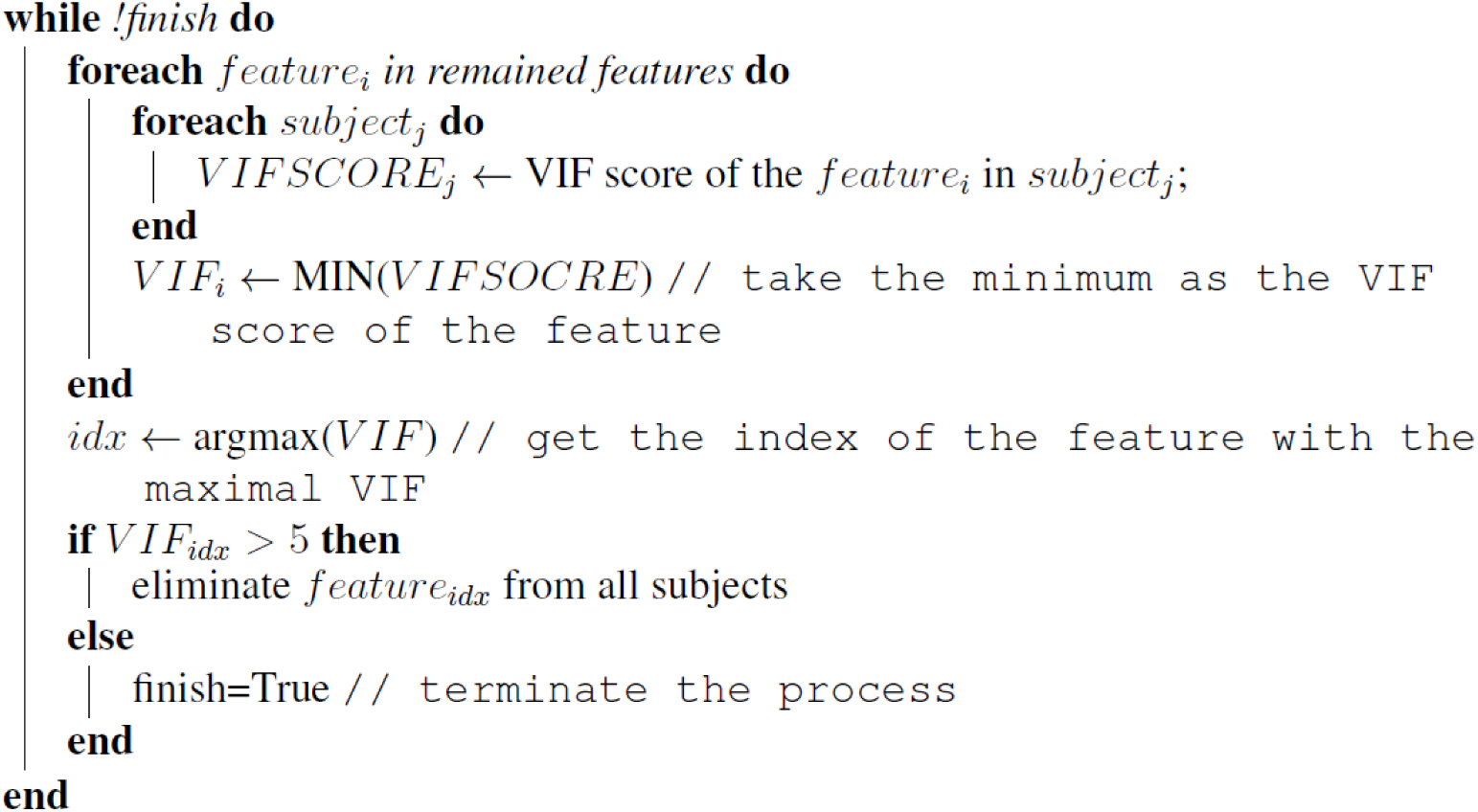
VIF-based multi-collinearity elimination pseudo code.

#### Models Evaluation

##### Classification categories

In experiment 1, there were two categories of classification: predicting the trial’s condition (whether sensorimotor conflict was introduced), or an objective classification, and predicting the subjective judgment, or a subjective classification. Each category is divided into a few sub-classifications.

In the objective category, there were five sub-classifications:

- Classifying trials as either congruent or incongruent.
- Classifying trials as either congruent or incongruent with 50ms delay.
- Classifying trials as either congruent or incongruent with 100ms delay.
- Classifying trials as either congruent or incongruent with 150ms delay.
- Classifying trials as either congruent or incongruent in the shuffled condition (see below).

In the *subjective* category, there were two sub-classifications:

- Classifying trials according to subjective judgment. Namely, predicting from the trial’s kinematics whether the participant judged the virtual movement as congruent with his real hand movement (had SoA over the movement) or not.
- Same, in shuffled condition (see below).

Experiment 2 included all of the abovementioned classification categories and an additional sub-classification in the objective category:

- Classifying trials to either congruent or incongruent with 200ms delay

##### Shuffled Condition

The two target labels in this study (sensorimotor conflicts and participant judgments) are correlated. As a consequence, the performance of the classifier in one category may be partially explained by the other target variable. For example, in the subjective classification, we are interested in how much information the hand kinematics holds regarding the subject’s answer. However, since the correlation between the magnitude of the sensorimotor conflict is highly correlated with the subject’s answer, there is a possibility that we would get high classification for the subject’s answer that is actually based on the sensorimotor conflict (via this correlation) and not the subject’s answer itself. The same applies when trying to classify the sensorimotor conflict, or the objective classification. To account for this relationship, we created the shuffled condition. The aim was to quantify how much of the classifier’s performance in one category is explained by the other target variable. This was accomplished by evaluating the classifier’s performance after shuffling the intended target labels while maintaining the correlation between them and the other target label. Thus, classifier performance measures solely the extent to which the other target variable explains classification performance and, as such, could serve as a new baseline for the normal condition. Comparing the normal and shuffled results (the shuffled gap) can reveal the extent to which classification performance depends on the intended target label or the other correlated target label.

To create the shuffled condition for the subjective classification, we randomly shuffled the subject’s answers (binary choice of yes/no) within each level of sensorimotor conflict. This reduced the link between the kinematics and subject’s answer, while preserving the correlation of the kinematic information to the sensorimotor conflict (the different magnitudes pf temporal delay). In this way we created a new baseline to which the normal condition can be compared. The gap between the shuffled and the normal conditions represents the information added solely by the subject’s answer.

To create the shuffled condition for the objective classification, we randomly shuffled the levels of the sensorimotor conflict (different magnitudes of temporal delay) within subjects’ answers (binary choice of yes/no). This reduced the link between the kinematics and sensorimotor conflict, while preserving the correlation of the kinematic information to the subject’s answers. In this way we created a new baseline to which the normal condition can be compared. The gap between the shuffled and the normal conditions represents the information added solely by the sensorimotor conflict.

##### Classifier Training and Evaluation

Classifiers were trained, using the full feature representation, for each classification in each participant separately. A 10-fold cross-validation method was used for the classification to evaluate classifier performance. Since the datasets for most classifications were imbalanced, AUC was employed as an evaluation metric. To determine the degree to which participants’ kinematic signatures are similar, the similarity between each participant’s logistic regression models was calculated. For each participant, a logistic regression model was trained using 10-fold cross-validation on the first objective classification for each fold using a clean feature representation, resulting in an overall of 10 models. Each participant’s representational model was the mean of the 10 model coefficients. Using the Pearson correlation between the coefficients of these models, the similarity between them was measured. The one-subject model technique was yet another method for demonstrating the similarities in the kinematic signatures of the participants. Instead of training a customized classifier for each participant, classifiers were trained iteratively on the data of a single participant and then evaluated on the data of the remaining participants.

##### Feature Importance Analysis

Both logistic regression and random forest feature importance analyses were extracted from models trained on a clean feature representation. The regression coefficients determined the importance of the features in the logistic regression models. The importance of the features in the random forest models was determined by calculating the mean decrease in impurity for each feature (Molnar, 2022).

#### Other Statistical Analyses

Participants’ performance was measured by quantifying the sensitivity and bias (Swets, 1964; Stanislaw and Todorov, 1999) of signal detection theory (SDT). Classifiers’ sensitivity and bias were also calculated to compare the performance of the classifiers to that of the participants. Pearson correlations were used to assess the relationship between model performance, participants’ SDT performance and questionnaire ratings. In addition to the shuffled condition, the Pearson correlation between the objective and subjective classifications was measured to account for the correlation between the objective-subjective performance gap (i.e., performance_objective_-performance_subjective_) and the participants’ sensitivity. In addition, the performance gap between the normal and shuffled conditions was calculated for both objective and subjective classifications (the objective shuffled gap and the subjective shuffled gap, respectively). Multiple comparisons were accounted for using the Bonferroni method.

## Results

### Clean Feature Representation Results

**Figure S3.**
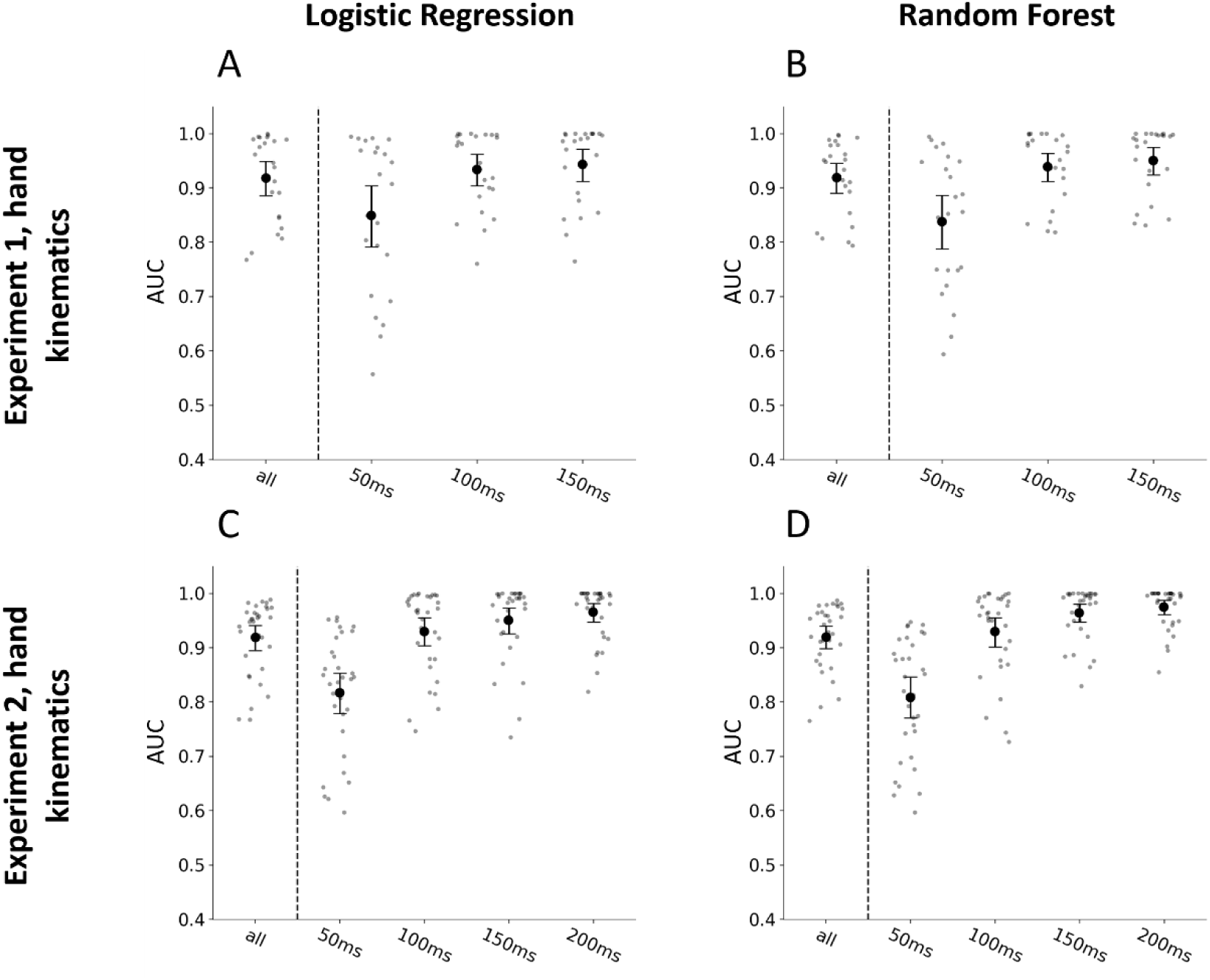
Classifier results using the *clean feature representation*. Average AUC across participants for each objective classification separately. The large black dot represents the group mean, the error bars represent the bootstrapped 95% confidence interval, and the small grey dots represent the individual participants. A, B: Experiment 1, hand kinematics (A) Logistic regression classifier (mean_all_=0.92, mean_50_=0.85, mean_100_=0.93, mean_150_=0.94) (B) Random forest classifier (mean_all_=0.92, mean_50_=0.84, mean_100_=0.94, mean_150_=0.95). C, D Experiment 2, hand kinematics. (C) Logistic regression classifier (mean_all_=0.92, mean_50_=0.81, mean_100_=0.93, mean_150_=0.95, mean_200_=0.97) (D) Random forest classifier (mean_all_=0.92, mean_50_=0.81, mean_100_=0.93, mean_150_=0.96, mean_200_=0.98).

### One Subject Model Performance

Another technique to demonstrate the similarities between the participants’ kinematic signatures was to, iteratively, train model on one subject and evaluate its performance on all of the other participants.

**Figure S4.**
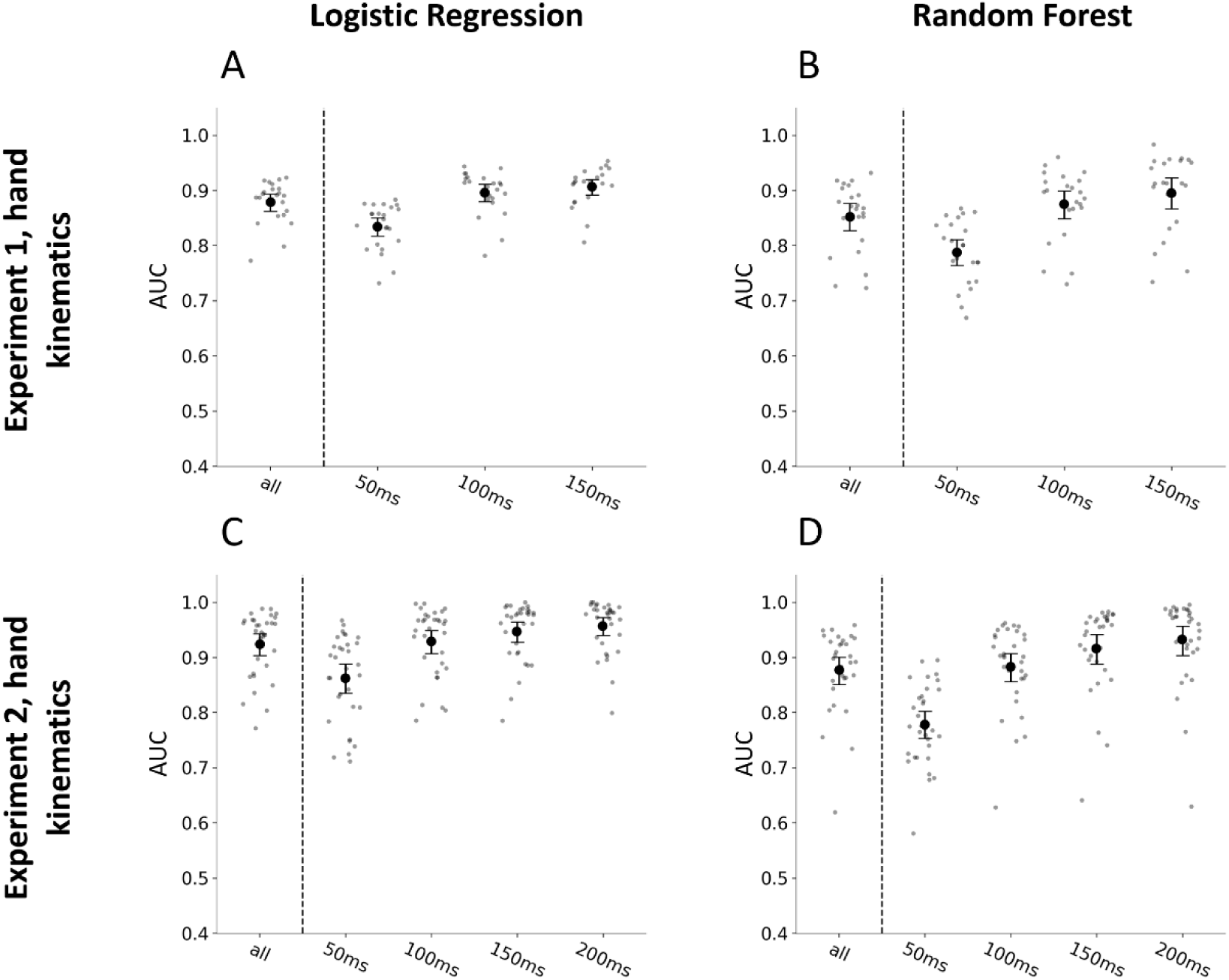
Classifier results using the *clean feature representation*. Average AUC across participants across each participant’s model for each objective classification separately. The model was trained on the first objective classification; therefore, all the trials have been used. The large black dot represents the group mean, the error bars represent the bootstrapped 95% confidence interval, and the small grey dots represent the individual participants’ mean across participants’ models. A, B: Experiment 1, hand kinematics (A) Logistic regression classifier (mean_all_=0.88, mean_50_=0.83, mean_100_=0.9, mean_150_=0.91) (B) Random forest classifier (mean_all_=0.85, mean_50_=0.79, mean_100_=0.88, mean_150_=0.9). C, D Experiment 2, hand kinematics. (C) Logistic regression classifier (mean_all_=0.92, mean_50_=0.86, mean_100_=0.93, mean_150_=0.95, mean_200_=0.96) (D) Random forest classifier (mean_all_=0.88, mean_50_=0.78, mean_100_=0.88, mean_150_=0.92, mean_200_=0.94).

### Subjective vs Objective Comparison

As both subjective and objective classification are related to the magnitude of the sensorimotor conflict we wish to see if the classification is more related to the sensory conflict or the subjective judgements. Thus, we computed the performance gap between the normal and shuffled conditions for both objective and subjective categories.

In experiment 1, the subjective shuffled gap was significant but relatively small (p<0.001 for both LR & RF) and significantly smaller than the reduction in classification when the objective data was shuffled gap (p<0.0001 for both LR & RF). (Figure S5 A&B). In addition, the Pearson correlation between the objective subjective performance gap (i.e., performance_objective_ - performance_subjective_) and the participants’ sensitivity was measured. There was a strong negative correlation between the objective subjective performance gaps and participants’ sensitivity (r_LR_=-0.67, p_LR_<0.001, r_RF_=-0.6, p_RF_=0.002). A high negative correlation indicates that the classifier’s performance on the subjective classification is heavily dependent on the kinematic signature of the sensorimotor conflict.

**Figure S5.**
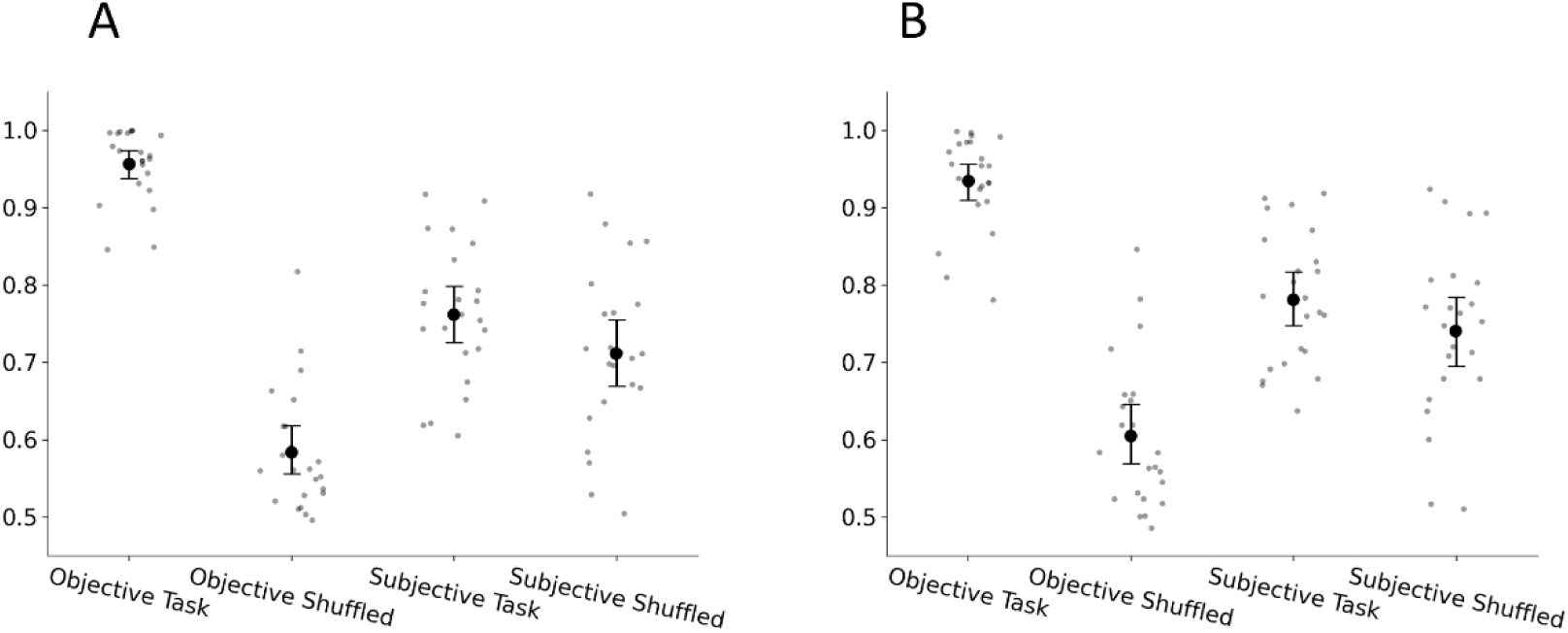
Classifier results of Experiment 1: subjective classification and shuffled condition. A,B average AUC across participants on both the objective and subjective classifications under normal and shuffled conditions. The large black dot represents the group mean, the error bars represent the bootstrapped 95% confidence interval, and the small grey dots represent individual participants (A) Logistic regression classifier (mean_objective_=0.96, mean_shuffled_objective_=0.58, mean_subjective_=0.76, mean_shuffled_subjective_=0.71) (B) Random forest classifier (mean_objective_=0.93, mean_shuffled_objective_=0.6, mean_subjective_=0.78, mean_shuffled_subjective_=0.74).

In experiment 2, the subjective shuffled gap was significant but relatively small (p<0.001 for both LR & RF) and significantly smaller than the objective shuffled gap (p<0.0001 for both LR & RF) (Figure S6 A&B). In addition, there was a strong negative correlation between the objective subjective performance gaps and participants’ sensitivity (r_LR_=-0.62, p_LR_<0.001, r_RF_=-0.59, p_RF_<0.001).

**Figure S6.**
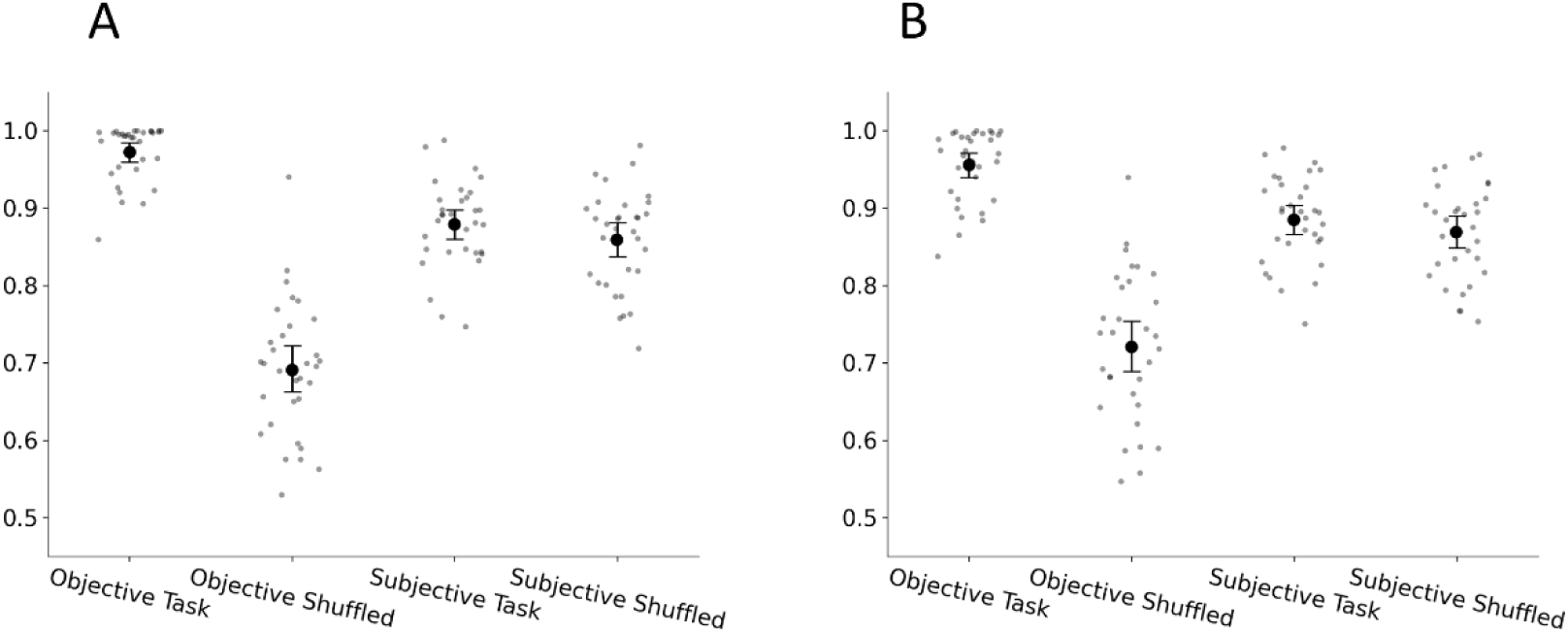
Classifier results of Experiment 2: subjective classification and shuffled condition. A,B average AUC across participants on both the objective and subjective classifications under normal and shuffled conditions. The large black dot represents the group mean, the error bars represent the bootstrapped 95% confidence interval, and the small grey dots represent the individual participants. (A) Logistic regression classifier (mean_objective_=0.97, mean_shuffled_objective_=0.69, mean_subjective_=0.87, mean_shuffled_subjective_=0.86) (B) Random forest classifier (mean_objective_=0.96, mean_shuffled_objective_=0.72, mean_subjective_=0.88,

